# CircRNA-based, non-integrated, *in vivo* panCAR-mediated B cell immune resetting in mouse models and non-human primates

**DOI:** 10.1101/2025.08.11.669448

**Authors:** Yanyan Wang, Xinyue Wang, Qian Pan, Haomeng Kou, Chi Zhang, Shengnan She, Jie Yin, Xinyuan Liao, Wanjia Wang, Ligaoyue Zhang, Huji Xu, Tianlei Ying, Yanling Wu, Liang Qu

## Abstract

B cell immune resetting, defined as the comprehensive depletion and subsequent reconstitution of B cells, represents a pivotal therapeutic approach for B cell-associated disorders. Recently, chimeric antigen receptor T cell (CAR-T) therapy has demonstrated substantial clinical efficacy in treating B cell malignancies and autoimmune diseases. However, this approach remains constrained by its reliance on lymphodepleting conditioning regimens, high invasiveness, high costs, and the risk of severe cytokine release syndrome (CRS). Antibody-based therapies, including monoclonal antibodies and T cell-engager (TCE) bispecific antibodies, suffer from suboptimal tissue penetration, resulting in inadequate B cell depletion in lymphoid and peripheral tissues, and subsequent rapid resurgence of pathogenic B cells. Here, we engineered highly stable circular RNAs (circRNAs) encoding anti-CD19 CAR (circRNA-CAR), which demonstrated prolonged and enhanced CAR protein expression compared to conventional linear mRNAs. By employing immune cell-tropic lipid nanoparticles (LNPs) for *in vivo* delivery, circRNA-CAR facilitated the generation of panCAR immune cells (including CAR-T, CAR-NK, and CAR-macrophages) in both murine models and cynomolgus monkeys. Compared to rituximab or teclistamab, *in vivo* panCAR induced more extensive B cell depletion across multiple tissue compartments in mice. Moreover, *in vivo* panCAR mediated robust B cell depletion, eliminated autoreactive antibodies, and restored renal function by alleviating proteinuria in a systemic lupus erythematosus (SLE) murine model. It also effectively reduced IgE antibody levels, decreased eosinophil activity, and improved lung pathology in an asthma murine model. Notably, *in vivo* panCAR-mediated B cell depletion significantly attenuated aging-associated phenotypes in 17- month-old aging mice, remarkably outperforming both rituximab and teclistamab. In cynomolgus monkeys, *in vivo* panCAR achieved sustained B cell depletion lasting approximately 40 to 50 days, followed by reconstitution of a naïve B cell repertoire. Collectively, circRNA-based *in vivo* panCAR enables complete B cell immune resetting, offering a versatile and promising therapeutic platform for a broad spectrum of B cell- and antibody-mediated disorders.

## INTRODUCTION

Pathogenic B cells and dysfunctional antibodies contribute not only to B cell lymphomas and autoimmune disorders but also to allergy, antibody-mediated aging, and neuropathies.^1–5^ Immunosuppressants are conventional pharmaceuticals employed in the treatment of autoimmune diseases; however, they are associated with substantial adverse effects and modest therapeutic benefits.^6–8^ B cell immune resetting, defined as the complete depletion of B cells followed by the subsequent reconstitution of B cell populations, provides a promising therapeutic approach for B cell- and antibody-associated diseases.^9^ While immunosuppressive medications require continuous administration to control autoimmune diseases, B cell immune resetting therapies offer the potential for fundamental and long-term remission.^10–12^ Antibody-based therapies, including monoclonal antibodies and T cell-engager (TCE) bispecific antibodies, have demonstrated efficacy in depleting B cells in peripheral blood but continue to suffer from suboptimal tissue penetration. ^13–17^ This limitation results in inadequate depletion of tissue-resident B cells in specific organs and subsequent rapid resurgence of pathogenic B cells, as observed in systemic lupus erythematosus (SLE).^13–17^

Chimeric antigen receptor T cell (CAR-T) therapies have demonstrated success in cancer treatment, with several studies supporting their feasibility for autoimmune diseases.^10,11,18–21^ The underlying rationale of CAR T cell therapy involves the use of B cell-targeting CAR-T cells to achieve profound B cell depletion. However, conventional autologous or allogeneic CAR-T therapies face significant translational barriers, including the requirement for lymphodepletion, high costs, risks of genotoxicity arising from integrating vectors, cytokine release syndrome (CRS), and neurotoxicity.^22–25^ Moreover, persistent CAR-T cell activity induces long-term B cell aplasia, severely impairing the reconstitution of humoral immunity.^26^

To overcome these constraints, off-the-shelf *in vivo* CAR-T platforms have emerged, offering streamlined manufacturing and reduced resource burdens.^27–29^ The utilization of DNA transposase-mediated or lentivirus-based *in vivo* CAR platforms carries the potential risk of permanent genomic integration.^30–32^ In addition, viral vectors can elicit anti-drug antibody (ADA) responses *in vivo*, thereby complicating repeated administration of virus-based *in vivo* CAR-T platforms.^33–35^ And the pre-existing antibodies can also directly neutralize these viral vectors *in vivo*, leading to decreased efficacy or complete ineffectiveness.^33–37^

Messenger RNAs (mRNAs) encapsulated within lipid nanoparticles (LNPs) to generate *in vivo* CAR-T cells have been pursued and have shown encouraging B cell depletion, as recently reported.^29,38^ mRNAs provide safety advantages through cytoplasmic protein translation without genomic integration. This approach additionally permits dose titration and repeated administration due to the properties of RNA lipid nanoparticle-based drugs.^39^ However, the inherent instability of mRNAs restricts the pharmacokinetic performance of mRNA-based drugs.^40^ In contrast, circular RNAs (circRNAs) are covalently closed circular RNA molecules that exhibit enhanced stability and expression capacity owing to their resistance to exonuclease degradation conferred by their ring structure.^41,42^ Therefore, circRNA-based *in vivo* CAR-T may enable more effective and durable production of CAR-expressing immune cells compared to mRNA-based approaches.

In addition to the RNA molecule encoding CAR proteins in the inner core, the LNP delivery vector encapsulated on the outer surface plays a crucial role in the efficacy of *in vivo* CAR.^43^ While cell-specific targeting by adding surface protein conjugators (e.g., exclusive T-cell transduction via anti-CD5/CD8 antibodies) has been established, such antibody-conjugated LNPs for *in vivo* targeted delivery often compromise stability, manufacturing yield, batch consistency, and storage feasibility.^44,45^ Recently, our group reported an *in vivo* panCAR platform that utilizes immune cell-tropic LNPs to deliver CAR-encoding circRNAs, thereby generating *in vivo* panCAR functional effector cells, including CAR-T, CAR-NK (natural killer) cells, and CAR-macrophages, demonstrating potent anti-tumor activity in preclinical models.^46^

Therefore, given the potential of circRNA-based *in vivo* panCAR for comprehensive B cell depletion and improved safety profiles, it is crucial to investigate whether this approach can achieve complete depletion of pathogenic B cells, including both circulating and tissue-resident populations. Moreover, this strategy holds promise as a versatile therapeutic platform for addressing diverse B cell- and antibody-mediated disorders beyond autoimmune diseases.

## RESULTS

### CircRNA translated chimeric antigen receptor (CAR) proteins and mediated effective cytotoxicity in human primary T cells and macrophages

We employed the Group I intron-based RNA circularization technique to generate circRNAs, encoding the anti-human-CD19 or anti-mouse-CD19 chimeric antigen receptor (CAR) proteins, referred to as circRNA^hCD19-CAR^ or circRNA^mCD19-CAR^ (Figure 1A).^42,47–49^ As a control, we also produced circRNA harboring only the circularization elements without the anti-CD19 CAR-encoding RNA sequence, referred to as circRNA^Ctrl^, which lacks the ability to encode proteins. Firstly, flow cytometry analysis demonstrated that both circRNA^hCD19-CAR^ and circRNA^mCD19-CAR^ efficiently expressed CAR proteins in HEK293T cells, and circRNA^hCD19-CAR^ also effectively encoded CAR proteins in human primary T cells (Figure 1B). To compare with linear mRNA, we also generated mRNA encoding the anti-human-CD19 or anti-mouse-CD19 CAR proteins, referred to as mRNA^hCD19-CAR^ or mRNA^mCD19-CAR^. These RNAs were transfected into HEK293T cells for subsequent assessment of CAR expression at different time points (Figure 1C). Flow cytometry results showed that both circRNA^hCD19-CAR^ and circRNA^mCD19-CAR^ achieved high-level CAR protein expression lasting approximately one week, while 1mΨ-modified mRNA^hCD19-CAR^ or mRNA^mCD19-CAR^ expressed CAR proteins for up to 3 days, and unmodified mRNA^hCD19-CAR^ or mRNA^mCD19-CAR^ produced CAR proteins lasting for almost 1 day (Figure 1C). These findings indicate the inherent high stability of circRNAs and the potential superiority of circRNA-based *in vivo* CAR.

**Figure 1.**
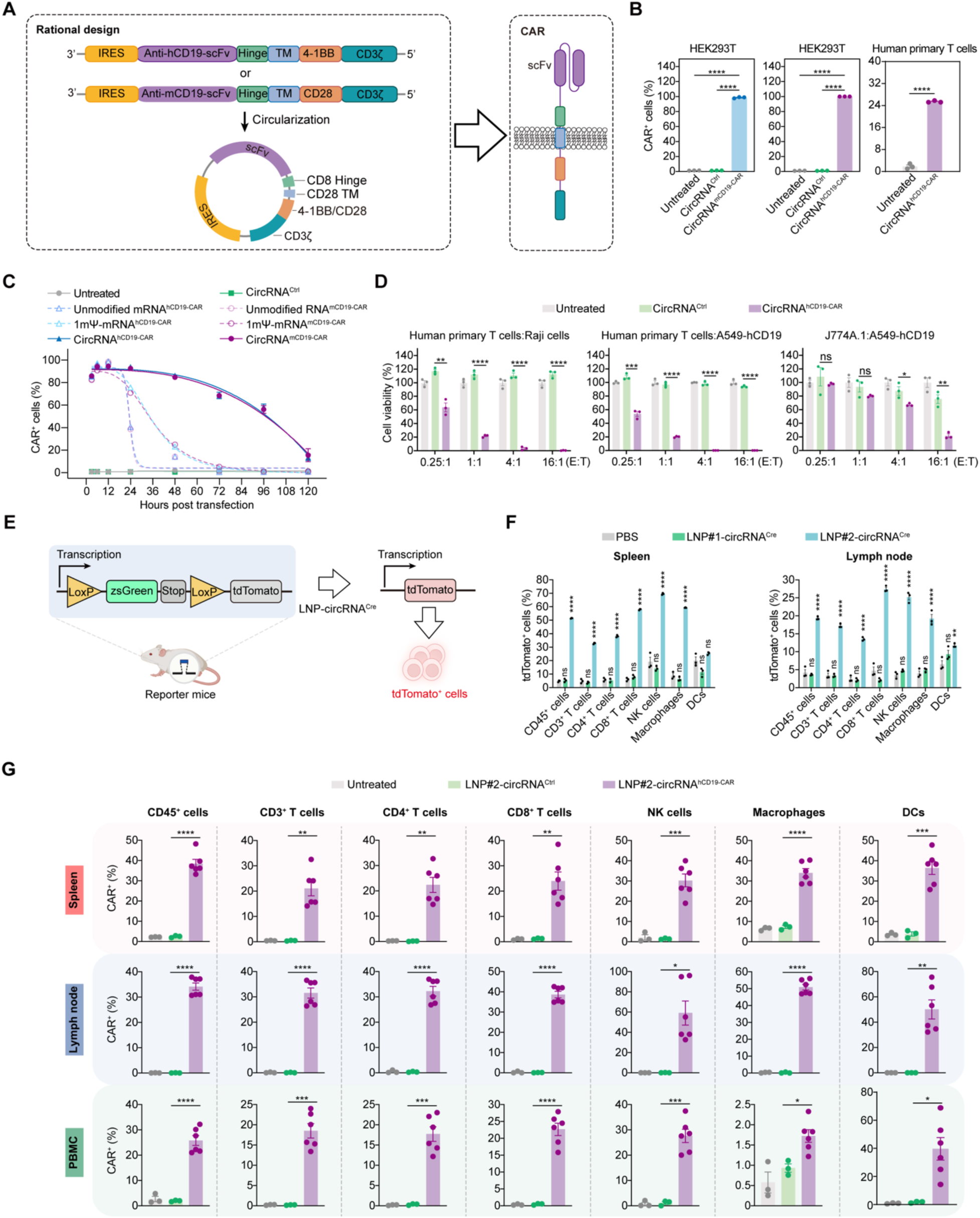
CircRNA mediated effective cytotoxicity in human primary T cells and generated *in vivo* panCAR via immunocyte-tropic LNP delivery in mice. (A) Schematic diagram illustrating the circularization of circRNA^hCD19-CAR^ and circRNA^mCD19-CAR^. (B) Detecting CAR protein expression in HEK293T cells and human primary T cells. (C) Comparative analysis of CAR expression levels in HEK293T cells following transfection with circRNA, 1mΨ-mRNA, or unmodified mRNA. (D) Human primary T cells and J774A.1 cells transfected with circRNA^hCD19-CAR^ exhibited cytotoxicity against A549-hCD19 and Raji cells. The cell viability was normalized with untreated group. (E) The schematic illustration shows that delivery of circRNA^Cre^ activates tdTomato expression in reporter mice via Cre-mediated genetic deletion of the stop cassette. (F) Evaluation of immune cell targeting efficiency of LNP#1-circRNA^Cre^ and LNP#2-circRNA^Cre^ in the spleen and lymph node of reporter mice using flow cytometry. (G) Detected CD19-CAR protein expression in multiple immune cell types within the spleen, lymph nodes and peripheral blood of mice intravenously administered with LNP#2-circRNA^hCD19-CAR^ using flow cytometry. In (B) to (G), data are presented as the mean ± SEM. In (B) to (G), an unpaired two-sided Student’s *t* test was conducted for comparison, as indicated. \**p* < 0.05; \*\**p* < 0.01; \*\*\**p* < 0.001; \*\*\*\**p* < 0.0001; ns, not significant. See also Figure S1 and S2.

Next, to investigate whether circRNA^hCD19-CAR^ could mediate specific cytotoxicity in human primary T cells or macrophages, we conducted cellular cytotoxicity assays by incubating circRNA^hCD19-CAR^-transfected human primary T cells or macrophages with Raji cells (CD19^+^ lymphoma cells) or hCD19-overexpressing A549 cells (A549-hCD19). The results showed that, compared to untreated or circRNA^Ctrl^ groups, circRNA^hCD19-CAR^-transfected human primary T cells or macrophages exhibited robust cytotoxic activity against Raji or A549-hCD19 cells in an effector:target (E:T) ratio-dependent manner (Figures 1D and S1A), indicating the potential of circRNA-based *in vivo* CAR therapy to mediate B cell depletion.

### Generation of *in vivo* panCAR via immunocyte-tropic lipid nanoparticles

Next, we employed immunocyte-tropic lipid nanoparticles (LNPs) to deliver circRNA^hCD19-CAR^ and circRNA^mCD19-CAR^, aiming to generate *in vivo* panCAR functional cell populations, including CAR-T cells, CAR-NK (natural killer) cells, and CAR-M (macrophage) cells.^46,50,51^ Two types of ionizable cationic lipids, LNP#1 and LNP#2, were used to encapsulated circRNAs, respectively. First, we used a Cre-reporter mouse model containing a CAG-loxp-zsGreen-stop-loxp-tdTomato-PolyA cassette integrated into the genome (Figure 1E). We generated circRNA^Cre^, encoding Cre recombinase, which was encapsulated in either LNP#1 or LNP#2, forming LNP#1-circRNA^Cre^ or LNP#2-circRNA^Cre^ nanoparticles. We demonstrated that LNP#2-circRNA^Cre^ induced tdTomato expression in T cells, macrophages, and NK cells in the spleen and lymph nodes, along with a notable decrease in zsGreen-positive cells across immune cell types, whereas LNP#1-circRNA^Cre^ failed to induce tdTomato expression in Cre-reporter mice (Figures 1F and S1B).

To further validate the *in vivo* panCAR formulation by circRNA, we delivered circRNA^hCD19-CAR^ or circRNA^Ctrl^ via LNP into mice. Six hours post intravenous injection of LNP-circRNA^hCD19-CAR^, peripheral blood, spleen, and lymph node samples were collected for flow cytometry analysis. Consistent with the Cre-reporter mouse results, LNP#2-circRNA^hCD19-CAR^ successfully expressed CAR proteins in various immune cell types, generating *in vivo* panCAR cell populations in spleen, lymph nodes, and peripheral blood (Figure 1G). Collectively, these results indicate that circRNA^hCD19-CAR^ can be delivered into multiple immune cell types to form *in vivo* panCAR via LNP#2 delivery.

### *In vivo* panCAR therapy achieved profound depletion of both circulating and tissue-resident B cells in mice

Next, we explored the effects of two LNP-encapsulated circRNA^mCD19-CAR^ formulations on B cell depletion in mice. The circRNA^mCD19-CAR^ was encapsulated with either LNP#1 or LNP#2 and intravenously injected into BALB/c mice twice at an interval of 3 days (Figure S2A). Flow cytometry results indicated that LNP#2-circRNA^mCD19-CAR^ achieved profound depletion of circulating B cells, whereas LNP#1-circRNA^mCD19-CAR^ exhibited minimal B cell depletion (Figures S2B-S2D). Therefore, LNP#2 was used to encapsulate circRNAs for *in vivo* delivery in subsequent experiments.

Previous studies have reported that teclistamab, a T cell-engager (TCE) bispecific antibody targeting CD3 and B cell maturation antigen (BCMA), mediates significant B cell depletion in clinical therapy for SLE patients.^15,16^ Rituximab, the approved anti-CD20 monoclonal antibody, is also used to deplete B cells for treating autoimmune diseases such as rheumatoid arthritis.^52–55^ However, Rituximab requires frequent high-dose administration and pre-treatment with analgesics or antihistamines.^56^ Another notable drawback of these antibody drugs is their limited efficacy against tissue-resident B cells, resulting in incomplete B cell depletion within lymphoid and peripheral tissues due to suboptimal tissue penetration.^57^ For systemic autoimmune diseases, profound depletion of tissue-resident B cells is crucial, otherwise, autoreactive B cells may rapidly expand, leading to disease relapse.

Therefore, we investigated whether circRNA-based *in vivo* panCAR could achieve complete and durable B cell depletion not only in peripheral blood but also in lymphoid and peripheral tissues (Figure 2A). We compared the effects of circRNA^hCD19-CAR^ with rituximab and teclistamab on B cell depletion in mice. BALB/c mice were intravenously administered circRNA^hCD19-CAR^, rituximab, or teclistamab four times at 3-day intervals. Subsequently, blood and organ samples were collected for flow cytometry, immunohistochemistry (IHC) staining and hematoxylin and eosin (H&E) staining analyses. Flow cytometry revealed that *in vivo* panCAR achieved complete depletion of CD19⁺ B cells, CD20⁺ B cells and CD138⁺ plasma cells in peripheral blood, whereas rituximab and teclistamab only modestly reduced B cell populations in this compartment (Figure 2B).

**Figure 2.**
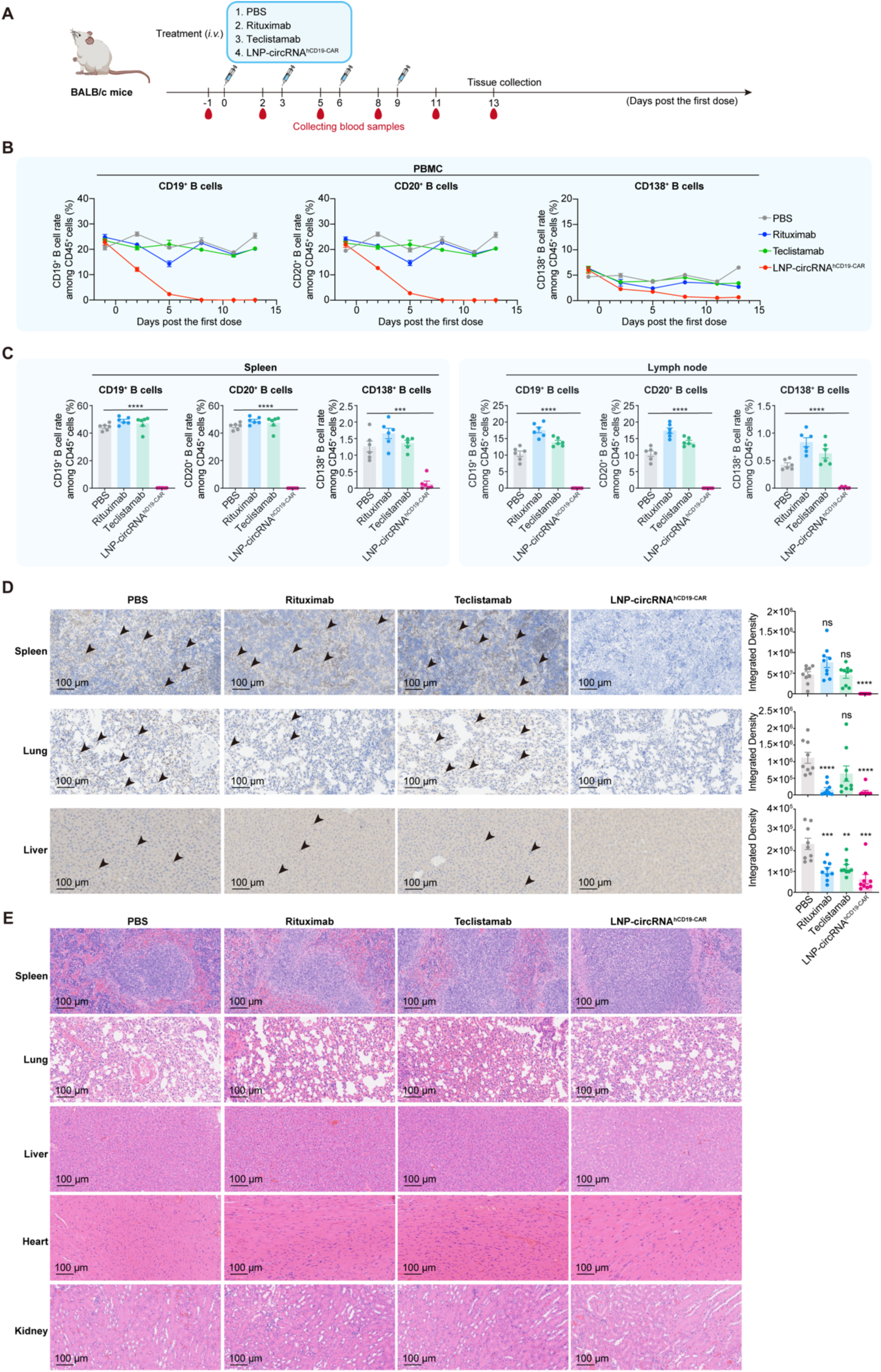
*In vivo* panCAR achieved profound depletion of both circulating and tissue-resident B cells in mice. (A) Schematic diagram depicting the administration and analysis of teclistamab, rituximab and circRNA^hCD19-CAR^ in wild-type mice. (B) Comparative effects of teclistamab, rituximab and *in vivo* panCAR on B cell depletion in peripheral blood. (C) Investigating the impact of distinct treatments on B cell depletion in the spleen and lymph nodes. Each symbol represents an individual mouse. (D) IHC staining was employed to assess the effects of teclistamab, rituximab and *in vivo* panCAR on tissue-resident CD19-positive B cells across various organs (liver, spleen, lung). The integrated density of IHC staining was quantified using Image J software. (E) H&E staining of various organs (liver, spleen, lung, heart and kidney) obtained from mice after PBS, teclistamab, rituximab or *in vivo* panCAR treatment. In (B) and (D), data are presented as the mean ± SEM. In (C) and (D), an unpaired two-sided Student’s *t* test was conducted for comparison, as indicated. \**p* < 0.05; \*\**p* < 0.01; \*\*\**p* < 0.001; \*\*\*\**p* < 0.0001; ns, not significant. See also Figure S3.

Furthermore, *in vivo* panCAR effectively depleted tissue-resident CD19⁺ B cells, CD20⁺ B cells and CD138⁺ plasma cells in both spleens and lymph nodes, while rituximab and teclistamab showed minimal efficacy in depleting tissue-resident B cells in these organs (Figure 2C). IHC staining confirmed significant B cell depletion in spleen, lung, and liver tissues in the panCAR-treated group compared to PBS, rituximab or teclistamab groups (Figure 2D). Notably, no CD19⁺ B cells were detected in the heart or kidneys of any treatment group (Figure S3).

To assess safety, H&E staining of major organs—including spleen, lung, liver, heart, and kidney—was performed. Similar to rituximab and teclistamab, *in vivo* panCAR treatment did not cause evident organ damage or abnormalities in these tissues (Figure 2E).

### *In vivo* panCAR achieved complete B cell depletion in systemic lupus erythematosus (SLE) autoimmune murine models

Next, we investigated whether *in vivo* panCAR-mediated complete B cell depletion could serve as an effective therapy for autoimmune disease. One hallmark of autoimmunity is the production of self-reactive B cells, which attack the body’s own tissues. Systemic lupus erythematosus (SLE) is an autoimmune disease characterized by antibody- and immune complex-mediated organ damage.^58^ Eliminating autoreactive B cells in both peripheral blood and tissues offers a promising strategy for achieving SLE remission and potential cure.^15^ MRL/MpJ-Fas^lpr^ mice, which carry a Fas^lpr^ mutation promoting survival of self-reactive lymphocytes, develop SLE-like symptoms with heterogeneity and complexity resembling human SLE.^49,59^ These mice were intravenously administered PBS, LNP-circRNA^mCD19-CAR^, or LNP-circRNA^hCD19-CAR^ five times consecutively at three-day intervals (Figure 3A). Flow cytometry analysis showed that both circRNA^mCD19-CAR^ and circRNA^hCD19-CAR^ induced complete depletion of CD19⁺ B cells, CD20⁺ B cells, and CD138⁺ plasma cells, while T cell populations remained unaffected (Figures 3B–3G, S4A and S4B).

**Figure 3.**
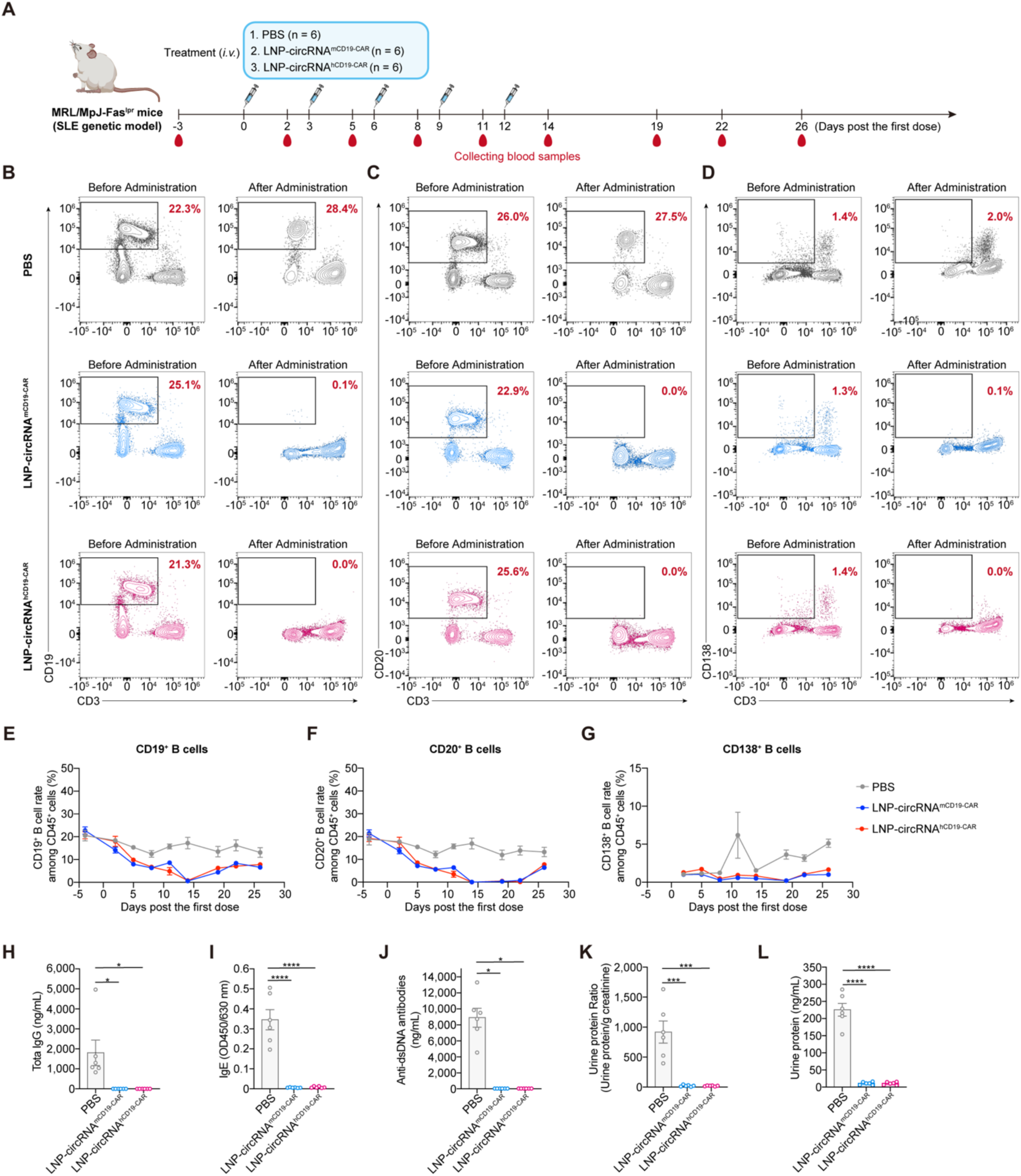
*In vivo* panCAR achieved complete B cell depletion in systemic lupus erythematosus (SLE) autoimmune murine models. (A) Schematic diagram depicting the administration and analysis of circRNA^mCD19-CAR^ or circRNA^hCD19-CAR^ in MRL/MpJ-Fas^lpr^ mice. (B-D) Flow cytometry plots showing CD19^+^ B cells(B), CD20^+^ B cells (C) and CD138^+^ B cells (D) depletion in peripheral blood of MRL/MpJ-Fas^lpr^ mice after circRNA^hCD19-CAR^ or circRNA^mCD19-CAR^ administration at 14 days post the first dose. (E-G) Effects of circRNA^hCD19-CAR^ or circRNA^mCD19-CAR^ on CD19^+^ B cells (E), CD20^+^ B cells (F) and CD138^+^ B cells (G) depletion in peripheral blood of MRL/MpJ-Fas^lpr^ mouse models. (H and I) Detecting the total IgG (H) and IgE (I) in the sera from treated mice via ELISA at 19 days post the first dose. (J) Detecting the anti-dsDNA antibody in the sera from treated mice via ELISA at 19 days post the first dose. (K and L) Detecting the ratios (urine protein to creatinine) (K) and urine protein (L) via ELISA at 19 days post the first dose. In (E) to (L), data are presented as the mean ± SEM. In (H) to (L), an unpaired two-sided Student’s *t* test was performed for comparison as indicated in the figures, \**p* < 0.05; \*\**p* < 0.01; \*\*\**p* < 0.001; \*\*\*\**p* < 0.0001; ns, not significant. Each symbol represents an individual mouse. See also Figure S4.

Following B cell depletion, sera and urine samples were collected from treated mice. ELISA assays revealed that both circRNA^mCD19-CAR^ and circRNA^hCD19-CAR^ treatments significantly reduced total IgG and IgE levels, as well as pathogenic anti-double-stranded DNA (anti-dsDNA) antibodies, compared to the PBS group (Figures 3H-3J). Proteinuria, a key clinical indicator of lupus nephritis, was markedly diminished in mice treated with either circRNA^mCD19-CAR^ or circRNA^hCD19-CAR^, indicating functional recovery and disease remission (Figures 3K and 3L).

Collectively, these findings demonstrate that *in vivo* panCAR-mediated B cell depletion and immune resetting effectively mitigate disease progression in systemic autoimmune mice, holding promise for achieving a cure in autoimmune disorders.

### *In vivo* panCAR significantly ameliorated the allergic asthma murine models

Asthma is one of the most prevalent chronic respiratory diseases, characterized by elevated IgE production, eosinophil infiltration, increased airway mucus secretion, and bronchial hyperresponsiveness.^10,60^ Eosinophil infiltration, driven by type 2 inflammatory cytokines (e.g., IL-4, IL-5 and IL-13) and B cells secreting specific antibodies, is a pivotal pathogenic factor in asthma.^60^ Consequently, drugs targeting IL-5/IL-5R and IL-4Rα have been approved for asthma treatment.^61^ CAR-T cell therapy strategies targeting eosinophils hold potential for achieving long-term asthma remission.^62,63^

In our previous results, we observed that *in vivo* panCAR reduced IgE levels in SLE mice (Figure 3I), inspiring us to investigate the potential effects of circRNA-based non-integrating panCAR on allergic asthma. Firstly, we established asthma mouse models following a previously reported protocol.^63^ BALB/c mice were sensitized via intraperitoneal injection of ovalbumin (OVA)/alum three times at 7-day interval, followed by daily intranasal OVA challenges. The asthma mice were randomly divided into two groups and intravenously administered circRNA^hCD19-CAR^ or PBS three times at 3-day interval (Figure 4A).

**Figure 4.**
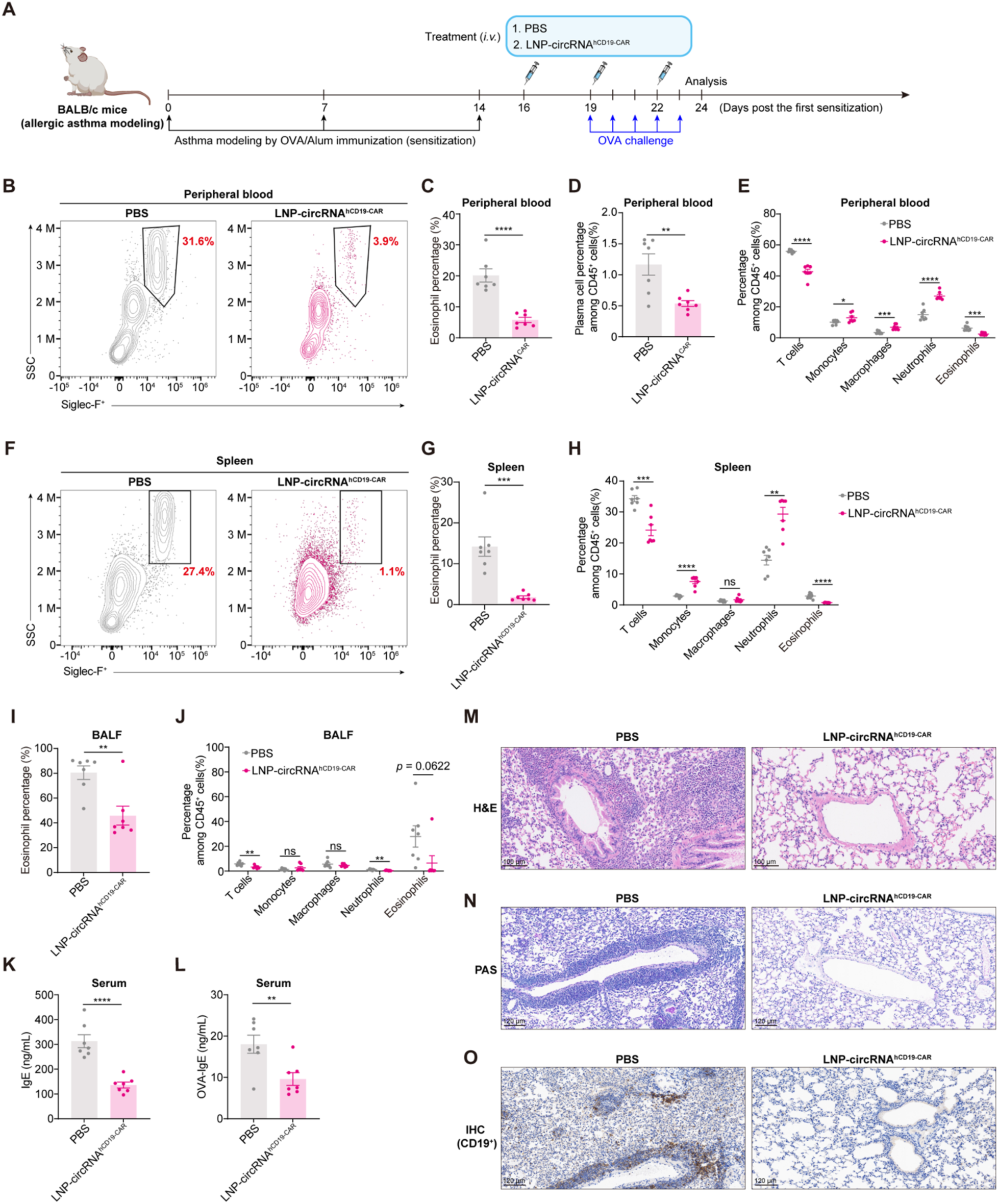
*In vivo* panCAR significantly ameliorated allergic asthma murine models. (A) Schematic diagram depicting the administration and analysis of circRNA^hCD19-CAR^ in OVA-induced asthma model of BALB/c mice. (B and C) Representative flow cytometry plots (B) and statistics (C) on eosinophils (Siglec-F^+^SSC^hi^) in peripheral blood. (D) Detecting on plasma cells in peripheral blood via flow cytometry. (E) Percentage of indicated immune cells in peripheral blood of mice with indicated treatment. T cells (CD3^+^); monocytes (CD11b^+^CD11c^-^Siglec-F^-^Ly6G^-^); macrophages (CD11b^+^CD11c^+^Siglec-F^+^); neutrophils (CD11b^+^Ly6G^+^); eosinophils (CD11b^+^CD11c^−^Siglec-F^+^). (F and G) Representative flow cytometry plots (F) and statistics (G) on eosinophils (Siglec-F^+^SSC^hi^) in spleen. (H) Percentage of indicated immune cells in spleen of asthma mice with indicated treatment. T cells (CD3^+^); monocytes (CD11b^+^CD11c^-^Siglec-F^-^Ly6G^-^); macrophages (CD11b^+^CD11c^+^Siglec-F^+^); neutrophils (CD11b^+^Ly6G^+^); eosinophils (CD11b^+^CD11c^−^Siglec-F^+^). (I) Detecting the eosinophils (Siglec-F^+^SSC^hi^) in BALF via flow cytometry. (J) Percentage of indicated immune cells in BALF of asthma mice with indicated treatment. T cells (CD3^+^); monocytes (CD11b^+^CD11c^-^Siglec-F^-^Ly6G^-^); macrophages (CD11b^+^CD11c^+^Siglec-F^+^); neutrophils (CD11b^+^Ly6G^+^); eosinophils (CD11b^+^CD11c^−^Siglec-F^+^). (K and L) ELISA examination of total IgE (K) and OVA-specific IgE (L) in the sera of mice with indicated treatment. (M) H&E staining was employed to assess the effects in lung tissues of asthma mice. (N) Periodic acid-Schiff (PAS) staining was used to analyze the level of mucus secretion in the lung tissues of asthmatic mice. (O) IHC staining showed the effect of CD19 positive cell after *in vivo* panCAR treatment in the lung tissues of asthma mice. In (C) to (L), data are presented as the mean ± SEM, and an unpaired two-sided Student’s *t* test was performed for comparison as indicated in the figures, \**p* < 0.05; \*\**p* < 0.01; \*\*\**p* < 0.001; \*\*\*\**p* < 0.0001; ns, not significant. Each symbol represents an individual mouse.

Flow cytometry analysis revealed a significant reduction in eosinophils and plasma cells, alongside minor increases in innate immune cells in peripheral blood (Figures 4B-4E). In the spleen, *in vivo* panCAR significantly reduced eosinophil proportions while increasing monocyte and neutrophil levels (Figures 4F-4H). Importantly, flow cytometry showed that *in vivo* panCAR markedly suppressed eosinophil and lymphocyte infiltration in bronchoalveolar lavage fluid (BALF) collected from treated asthma mice (Figures 4I and 4J). Serum levels of total IgE and OVA-specific IgE were also significantly lower in panCAR-treated mice compared to PBS controls (Figures 4K and 4L).

Moreover, hematoxylin and eosin (H&E) staining and periodic acid–Schiff (PAS) staining demonstrated that *in vivo* panCAR reduced inflammatory cell infiltration and mucus secretion in alveoli and bronchi of lung tissues (Figures 4M and 4N). Additionally, compared to PBS, *in vivo* panCAR treatment resulted in significant depletion of CD19⁺ B cells in lung tissues (Figure 4O).

Collectively, these results demonstrate that *in vivo* panCAR effectively ameliorates asthma symptoms in mouse models, providing potential therapeutic insights for B cell- and IgE-associated allergic diseases.

### *In vivo* panCAR attenuated aging-associated phenotypes in 17-month-old aging mice

Recent research has revealed elevated immunoglobulin accumulation at senescence-sensitive regions (SSRs), contributing to multi-organ senescence in mammals.^64^ Over-accumulation of IgG in various organs promotes chronic inflammation, thereby exacerbating tissue degeneration.^64^ Aging is a natural, complex, and progressive decline in functional integrity and physiological processes across tissues over time, ultimately resulting in death.^65^ Chronic inflammation, recognized as one of the fourteen hallmarks of aging, manifests as persistently elevated systemic inflammation and is strongly linked to various aging-related pathologies, including atherosclerosis, neuroinflammation, osteoarthritis, and intervertebral disc degeneration.^66^ Critically, studies have established that immune system aging drives the aging of solid organs in mammals.^67^ Consequently, the immune system,especially B cell responses and antibodies,represents a fundamental driver of aging.

The above results demonstrate that *in vivo* panCAR enables profound depletion of both circulating and tissue-resident B cells in multiple organs, as well as effectively decreases total IgG levels (Figures 2 and 3). Therefore, we further investigated whether *in vivo* panCAR could ameliorate aging-associated phenotypes by mediating complete B cell depletion in aging mouse models (Figure 5A). Seventeen-month-old BALB/c mice were intravenously administered PBS, rituximab, teclistamab, or circRNA^hCD19-CAR^ four times at 3-day interval. Subsequently, blood and organ tissue samples were collected for subsequent analysis. Consistent with previous findings, flow cytometry analysis showed that *in vivo* panCAR achieved complete depletion of circulating CD19⁺, CD20⁺ and CD138⁺ B cells in aging mice, whereas rituximab or teclistamab only achieved a slight reduction in B cell populations in peripheral blood compared to the PBS group (Figures 5B and 5C). Notably, we also found *in vivo* panCAR exhibited slight increases in innate immune cells such as macrophages and NK cells (Figure S5A).

**Figure 5.**
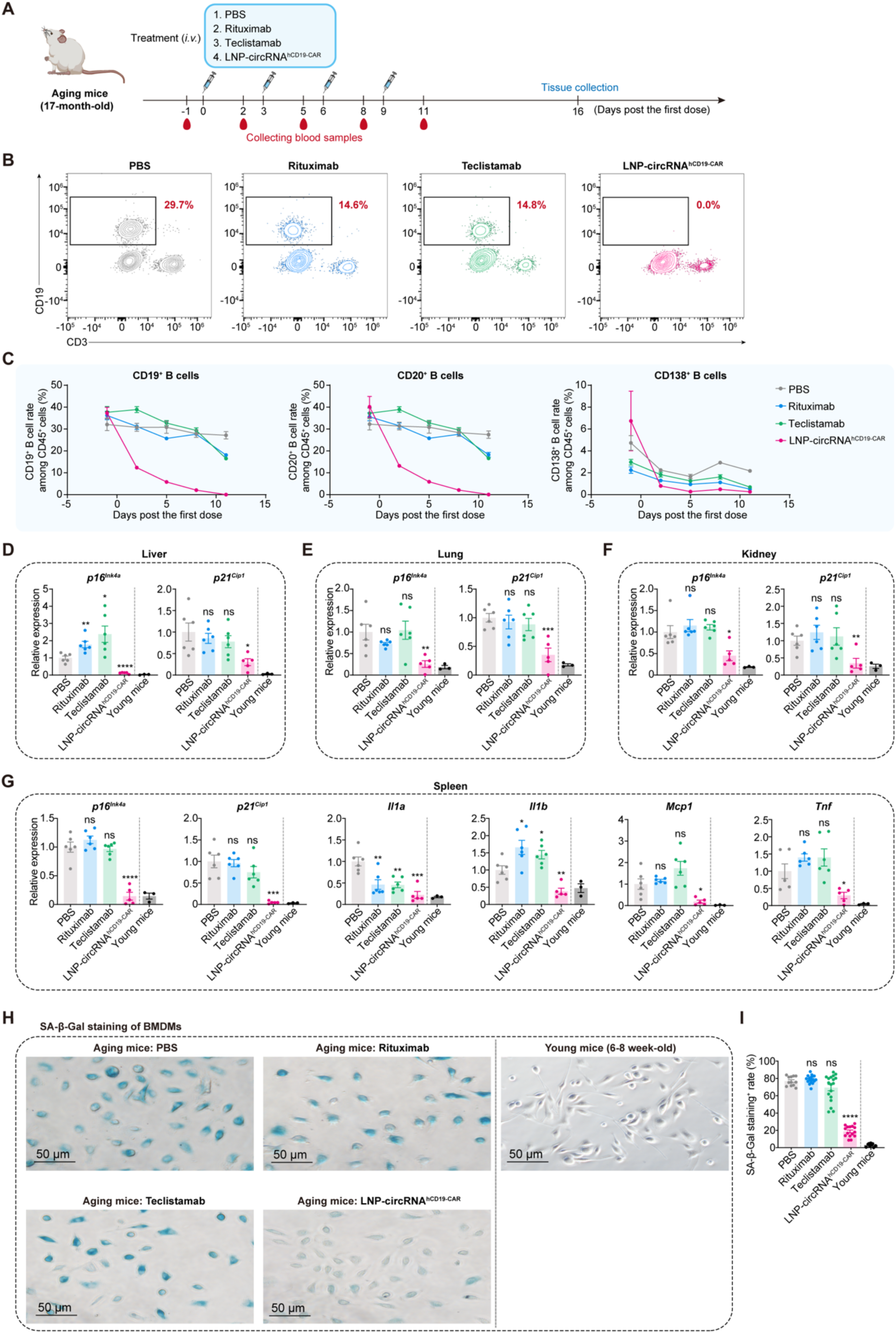
*In vivo* panCAR attenuated aging-associated phenotypes in 17-month-old aging mice. (A) Schematic diagram depicting the administration of PBS, rituximab, teclistamab or circRNA^hCD19-CAR^ in aging mice. (B) Flow cytometry plots on CD19^+^ B cell depletion at 48 hours post the final administration. (C) Comparation of *in vivo* panCAR with antibody-based therapeutics on B cells in the peripheral blood of senescent mice. (D-F) Comparative analysis on senescence-associated gene expression in liver (D), lung (E) and kidney (F) using RT-qPCR. Each symbol represents an individual mouse. (G) SASP analysis in spleen by RT-qPCR across treatment groups. Each symbol represents an individual mouse. (H) SA-β-Gal staining analysis of BMDMs from young (6-8-week-old) and aging (17-month-old) mice at 16 days post the first administration. (I) The SA-β-Gal positive cells in (H) was quantified using ImageJ software. In (C) to (G) and (I), data are presented as the mean ± SEM, and an unpaired two-sided Student’s *t* test was performed for comparison as indicated in the figures, \**p* < 0.05; \*\**p* < 0.01; \*\*\**p* < 0.001; \*\*\*\**p* < 0.0001; ns, not significant. See also Figures S5.

To further explore the effects of *in vivo* panCAR-mediated B cell depletion on aging-associated phenotypes, major organs (including heart, liver, spleen, lung and kidney) from treated aging mice were collected to analyze senescence-associated markers. Quantitative PCR revealed that *in vivo* panCAR significantly suppressed the levels of key aging-related hallmarks (*p16^Ink4a^*, *p21^Cip1^*) across all analyzed tissues, whereas the teclistamab and rituximab groups showed minimal effects (Figures 5D-5G and S5B). Additionally, we assessed the senescence-associated secretory phenotype (SASP) using spleen samples. *In vivo* panCAR significantly downregulated the expression of SASP factors (e.g., *Il1b*, *Mcp1*) compared to the PBS group, while teclistamab and rituximab exhibited minimal effects on reducing SASP-related genes expression (Figure 5G).

The activity of β-galactosidase (β-Gal) is an important biomarker of senescence in the cellular biology of aging, and senescence-associated β-galactosidase (SA-β-Gal) can be used to detect and analyze the presence of senescent cells.^68^ Therefore, we isolated bone marrow-derived macrophages (BMDMs) and assessed cellular senescence using SA-β-Gal staining.^69^ Morphologically, the BMDMs from aging mice treated with PBS exhibited a flattened morphology and abnormal enlargement, and SA-β-Gal staining confirmed a significantly increased number of senescent cells in aging mice compared to 6-8-week-old young mice (Figure 5H). The *in vivo* panCAR group significantly reduced the proportion of SA-β-Gal-positive BMDMs, while the teclistamab group showed only minimal effects, and the rituximab group demonstrated almost no detectable effect (Figure 5H and 5I). In summary, our results indicate that *in vivo* panCAR effectively mitigates aging-related phenotypes in aging mice.

### *In vivo* panCAR achieved complete B cell depletion in non-human primates (NHPs)

To further evaluate the efficacy and safety of circRNA-based *in vivo* panCAR in non-human primates (NHPs), we subsequently performed experiments in cynomolgus monkeys. Two monkeys, aged 2 to 4 years, received intravenous administration of LNP-circRNA^hCD19-CAR^ on days 0, 2, and 6, followed by peripheral blood collection at various time points for analysis (Figure 6A). Consistent with murine studies, we found that circRNA^hCD19-CAR^ was delivered into CD3⁺ T cells, macrophages and NK cells and expressed hCD19-CAR proteins in various CD45⁺ immune cell types for nearly two weeks, indicating that LNP-circRNA^hCD19-CAR^ generated *in vivo* panCAR (CAR-T, CAR-NK, and CAR-macrophage) cells in cynomolgus monkeys (Figures 6B-6E).

**Figure 6.**
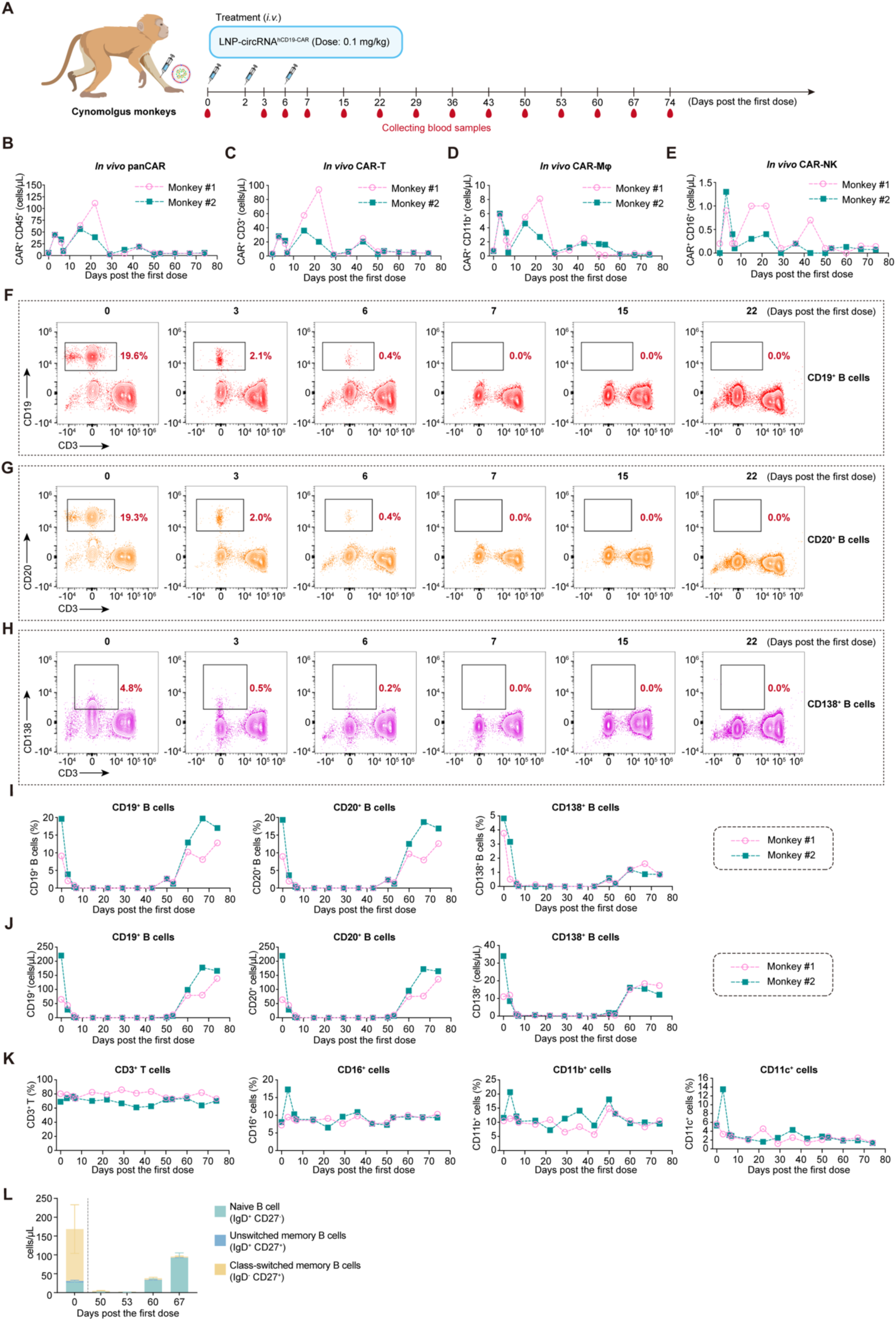
*In vivo* panCAR achieved complete B cell depletion in non-human primates (NHPs) (A) Schematic diagram of circRNA^hCD19-CAR^ administration in cynomolgus monkeys. (B-E) Evaluating the expression of *in vivo* panCAR on CD45+ immune cells (B), CD3^+^ T cells (C), CD11b^+^ cells (D) or CD16^+^ cells (E), in peripheral blood of cynomolgus monkey. (F-H) Flow cytometry plots on CD19^+^ B cells (F), CD20^+^ B cells (G) and CD138^+^ B cells (H) depletion in cynomolgus monkeys. (I and J) Percentage (I) or cell count quantification (J) of CD19^+^ B cells, CD20^+^ B cells and CD138^+^ B cells in peripheral blood of cynomolgus monkey treated with *in vivo* panCAR at different time points. (K) Percentage of CD3^+^ T cells, CD16^+^ cells, CD11b^+^ cells and CD11c^+^ cells in peripheral blood of cynomolgus monkey treated with *in vivo* panCAR at different time points. (L) Detecting the subtypes of B cell populations before treatment or after treatment of *in vivo* panCAR. In (L), data are presented as the mean ± SEM. See also Figure S6.

Next, flow cytometry results demonstrated that circRNA-based *in vivo* panCAR achieved complete depletion of CD19⁺ B cells, CD20⁺ B cells and CD138⁺ plasma cells in cynomolgus monkeys, with profound B cell depletion lasting approximately 40 to 50 days (Figures 6F-6J). In contrast, *in vivo* panCAR exhibited minimal effects on CD3⁺ T cells, CD16^+^ cells, CD11b^+^ cells, and CD11c^+^ cells, underscoring the high specificity of panCAR-mediated B cell depletion in cynomolgus monkeys (Figures 6K and S6).

Importantly, we also observed the subsequent reconstitution of B cells in the peripheral blood of cynomolgus monkeys after approximately 40-50 days of B cell depletion mediated by *in vivo* panCAR. The major component of newly regenerated B cells consisted mainly of IgD⁺ CD27⁻ naïve B cells. In summary, these results indicate that circRNA^hCD19-CAR^ can generate *in vivo* panCAR cells and induce B cell immune resetting in non-human primates (Figure 6L), establishing a foundation for clinical translation potential in the treatment of autoimmune diseases and other B cell- and antibody-associated disorders.

### Safety evaluation of *in vivo* panCAR in non-human primates (NHPs)

To further evaluate the safety of circRNA-based *in vivo* panCAR in NHPs, we collected serum from cynomolgus monkeys treated with *in vivo* panCAR. ELISA results showed that pro-inflammatory cytokines, including IL-6, TNF-α, IL-1β and monocyte chemoattractant protein-1 (MCP-1), exhibited no significant elevation after treatment with *in vivo* panCAR, fluctuating only

within a very narrow range (Figures 7A-7D). Concurrently, we further investigated whether *in vivo* panCAR resulted in potential hepatotoxicity in NHPs. We detected three classic biomarkers of hepatotoxicity, including alanine aminotransferase (ALT), aspartate aminotransferase (AST) and alkaline phosphatase (ALP), as well as C-reactive protein (CRP). The results revealed mild elevations in all three aminotransferases and CRP after administration of *in vivo* panCAR, with levels of the aminotransferases promptly returning to baseline (Figures 7E-7H). We speculate that this transient effect might be attributable to the potential immunogenicity of circRNA or the adjuvant activity of LNPs.^70–74^ Collectively, these results provide preliminary proof of safety for circRNA-based *in vivo* panCAR in NHPs, indicating its potential for clinical translation.

**Figure 7.**
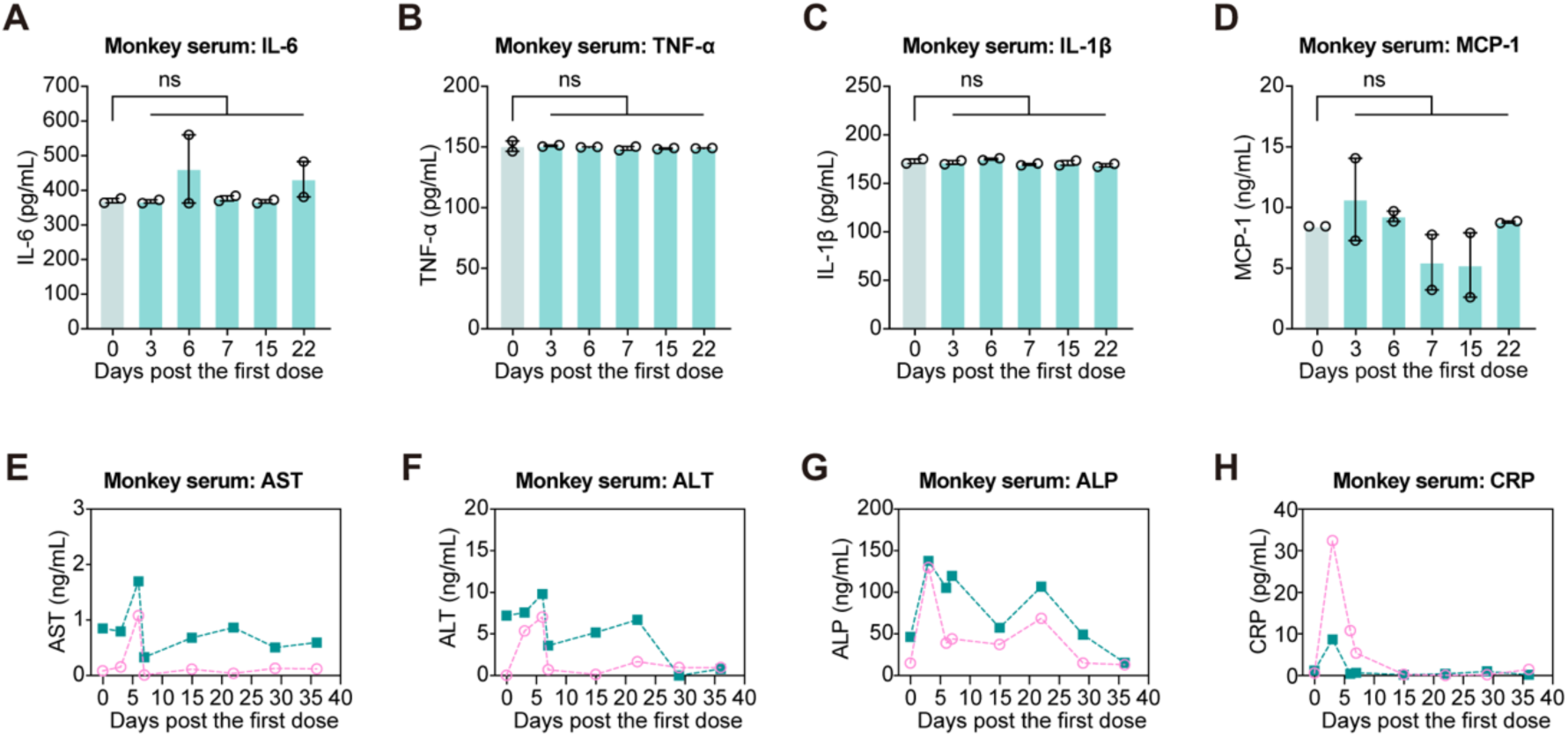
Safety evaluation of *in vivo* panCAR in non-human primates (NHPs) (A-D) Evaluating the effects of *in vivo* panCAR on pro-inflammatory cytokines including IL-6 (A), TNF-α (B), IL-1β (C) and MCP-1 (D) in the sera of treated cynomolgus monkey using ELISA. (E-G) Evaluating the effects of *in vivo* panCAR on hepatotoxicity indicators including AST (E), ALT (F) and ALP (G) in the sera of treated cynomolgus monkey using ELISA. (H) Evaluating the effects of *in vivo* panCAR on C-reactive protein (CRP) in the sera of treated cynomolgus monkey using ELISA. Data are presented as the mean ± SEM. In (A) to (D), an unpaired two-sided Student’s *t* test was performed for comparison as indicated in the figures, \**p* < 0.05; \*\**p* < 0.01; \*\*\**p* < 0.001; \*\*\*\**p* < 0.0001; ns, not significant.

## DISCUSSION

Autoimmune diseases (AIDs), such as systemic lupus erythematosus (SLE) and rheumatoid arthritis, are primarily characterized by the activation of autoreactive B cells, resulting in multi-organ damage.^75,76^ B cells play a pivotal role in the development of these disorders, particularly through the production of autoreactive antibodies which mediate tissue injury.^77,78^ Consequently, targeted B cell depletion has emerged as a critical therapeutic strategy for treating AIDs.^10,15,19–21,29,77,79^ While rituximab effectively depletes B cells in peripheral blood, it often fails to achieve complete depletion of B cells within tissues and rarely induces sustained drug-free remission, often leading to rapid disease relapse.^14^ T cell engagers (TCEs) have demonstrated applicability in treating AIDs through B cell depletion; however, achieving profound depletion of tissue-resident B cells within organs remains challenging.^15–17^ This limitation likely results from TCEs’ dependence on effector T cell proximity and the limited tissue penetration of antibody drugs.^15–17^

B cell-targeted chimeric antigen receptor T (CAR-T) cell therapy has been recognized as an innovative therapeutic approach, demonstrating potential for inducing long-term drug-free remission through deep B cell depletion.^10,19,49,79^ However, both autologous and allogeneic CAR-T applications are associated with challenges, including reliance on lymphodepleting conditioning regimens, the high invasiveness of procedures, high costs, safety concerns regarding genomic integration, cytokine release syndrome (CRS) and neurotoxicity.^80^ The *in vivo* CAR strategy, although previously used primarily in ex vivo gene therapy, employs genome-integrative lentiviral vectors that not only have the potential to cause genomic off-target effects due to random integration into genomic DNA but may also induce anti-drug antibody (ADA) production, thereby preventing administration of repeat dosing upon disease relapse.^81,82^

In this study, we introduced a circRNA-based non-integrated *in vivo* panCAR strategy, which not only efficiently depletes B cells in both peripheral blood and tissues/organs but also subsequently resets the B cell populations, providing a potential avenue for the treatment of autoimmune diseases. Due to its distinct circular structure, circRNA is highly stable and resistant to exonuclease-mediated degradation, thereby ensuring prolonged CAR expression and sustained immune cell clearance *in vivo* (Figure 1B). Compared to antibody-based therapies such as teclistamab and rituximab, *in vivo* panCAR achieves complete depletion of the circulating B cells in peripheral blood and tissue-resident B cells within organs (Figure 2). Moreover, we demonstrated proof-of-concept of circRNA-based *in vivo* panCAR in treating SLE, allergic asthma, and attenuating aging-associated phenotypes in 17-month-old aging mice (Figures 3-5). Validation in non-human primates further demonstrated that *in vivo* panCAR achieves comprehensive B cell depletion in NHPs with a sustained B cell depletion state lasting over one month, along with subsequent immune resetting of the B cell population, primarily consisting of naïve B cells (Figure 6). These findings emphasize the therapeutic potential of *in vivo* panCAR in achieving long-term remission following drug-free treatment of autoimmune diseases.

Indeed, B cells represent a pivotal element of the adaptive immune response, contributing to the maintenance of immune homeostasis within the body and exhibiting a strong association with various diseases. In principle, circRNA-based *in vivo* panCAR-mediated B cell depletion is potentially applicable to the treatment of most diseases associated with B cell abnormalities. This study uses autoimmune diseases, allergic diseases, and immune aging as representative models in a proof-of-concept investigation. In the future, the *in vivo* panCAR-mediated B cell immune resetting tool may provide a promising therapeutic platform for a broad spectrum of B cell- and antibody-related disorders beyond autoimmune diseases.

---

**Figure S1.**
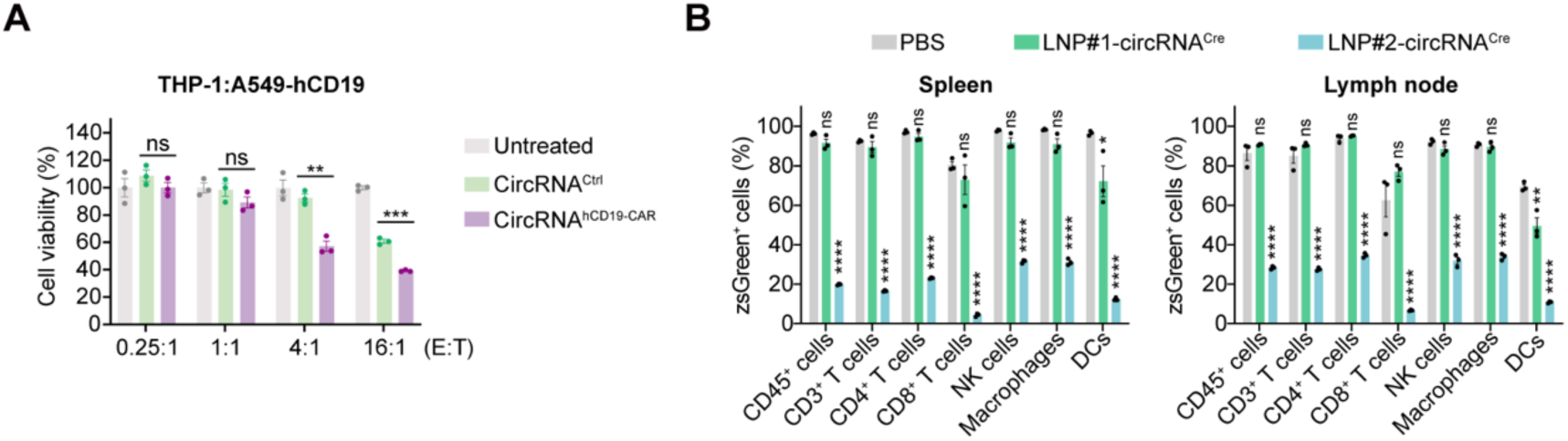
CircRNA mediated effective cytotoxicity in macrophages and generated *in vivo* panCAR via immunocyte-tropic LNP delivery in mice, related to Figure 1. (A) THP-1 cells transfected with circRNA^hCD19-CAR^ exhibited cytotoxicity against A549-hCD19 tumor cells. (B) Assessing the effects of circRNA^Cre^ on the proportion of zsGreen-positive cells among different immune cell populations in the spleen and lymph nodes. In (A) and (B), data are presented as the mean ± SEM, and an unpaired two-sided Student’s *t* test was performed for comparison as indicated in the figures, \**p* < 0.05; \*\**p* < 0.01; \*\*\**p* < 0.001; \*\*\*\**p* < 0.0001; ns, not significant.

**Figure S2.**
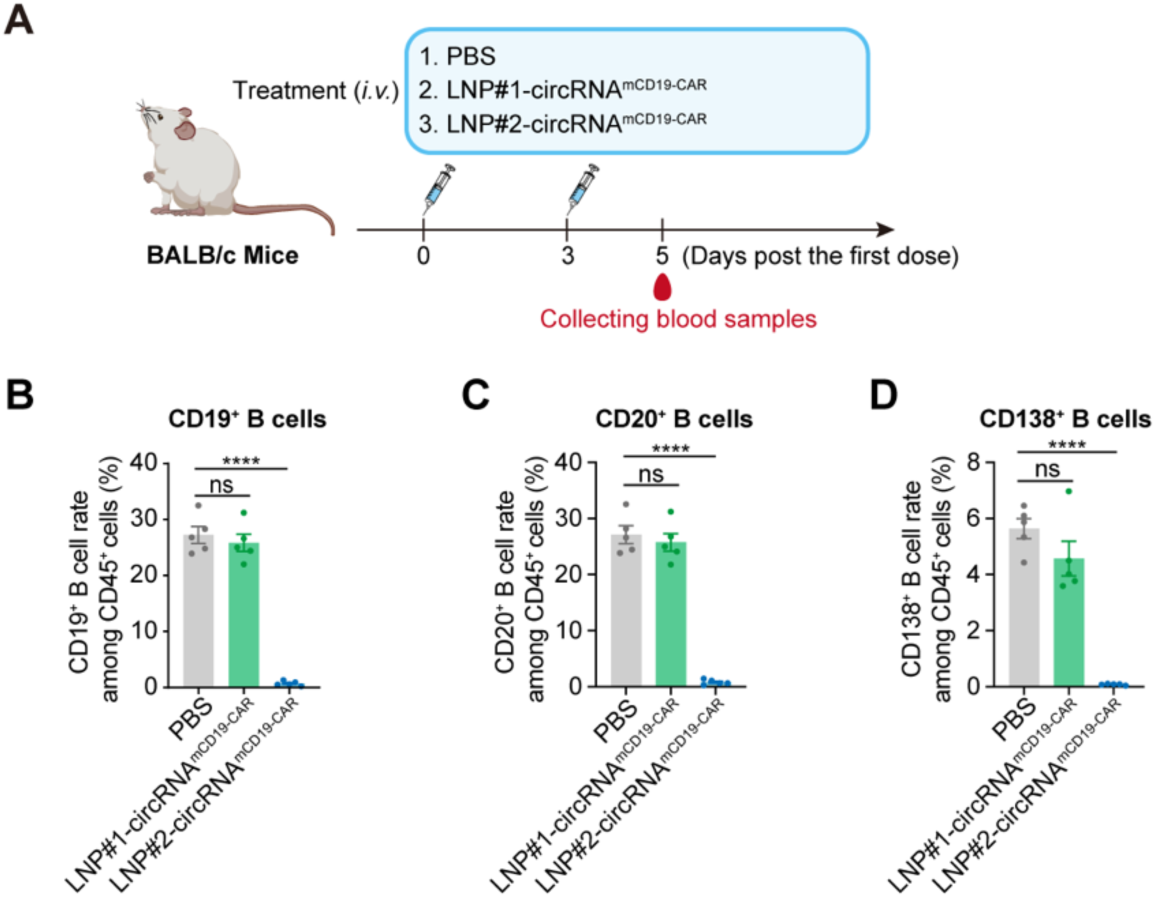
CircRNA mediated complete B cells depletion via immunocyte-tropic lipid nanoparticles (LNPs) delivery in mice, related to Figure 1. (A) Schematic diagram depicting the administration and analysis of LNP#1-circRNA^mCD19-CAR^ and LNP#2-circRNA^mCD19-CAR^ in wild-type mice. (B-D) Effects of circRNA^mCD19-CAR^ on CD19^+^ B cells (B), CD20^+^ B cells (C) and CD138^+^ B cells (D) depletion in the peripheral blood of treated mice at 5 days post the first dose. In (B) to (D), data are presented as the mean ± SEM, and an unpaired two-sided Student’s *t* test was performed for comparison as indicated in the figures, \**p* < 0.05; \*\**p* < 0.01; \*\*\**p* < 0.001; \*\*\*\**p* < 0.0001; ns, not significant.

**Figure S3.**
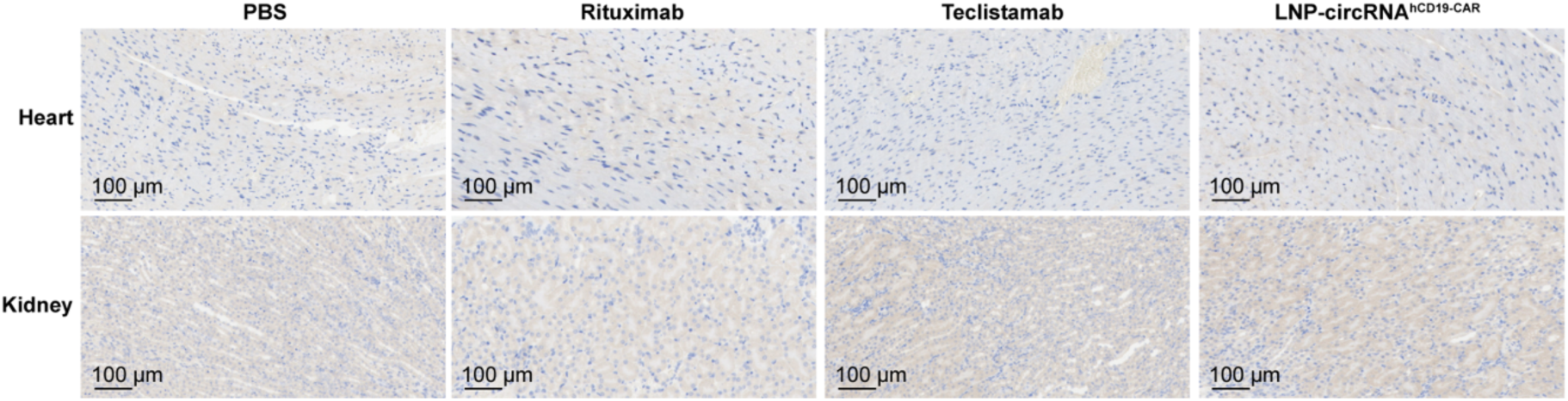
IHC staining (anti-CD19) using the heart or kidney tissues from mice treated with PBS, teclistamab, rituximab or *in vivo* panCAR, related to Figure 2.

**Figure S4.**
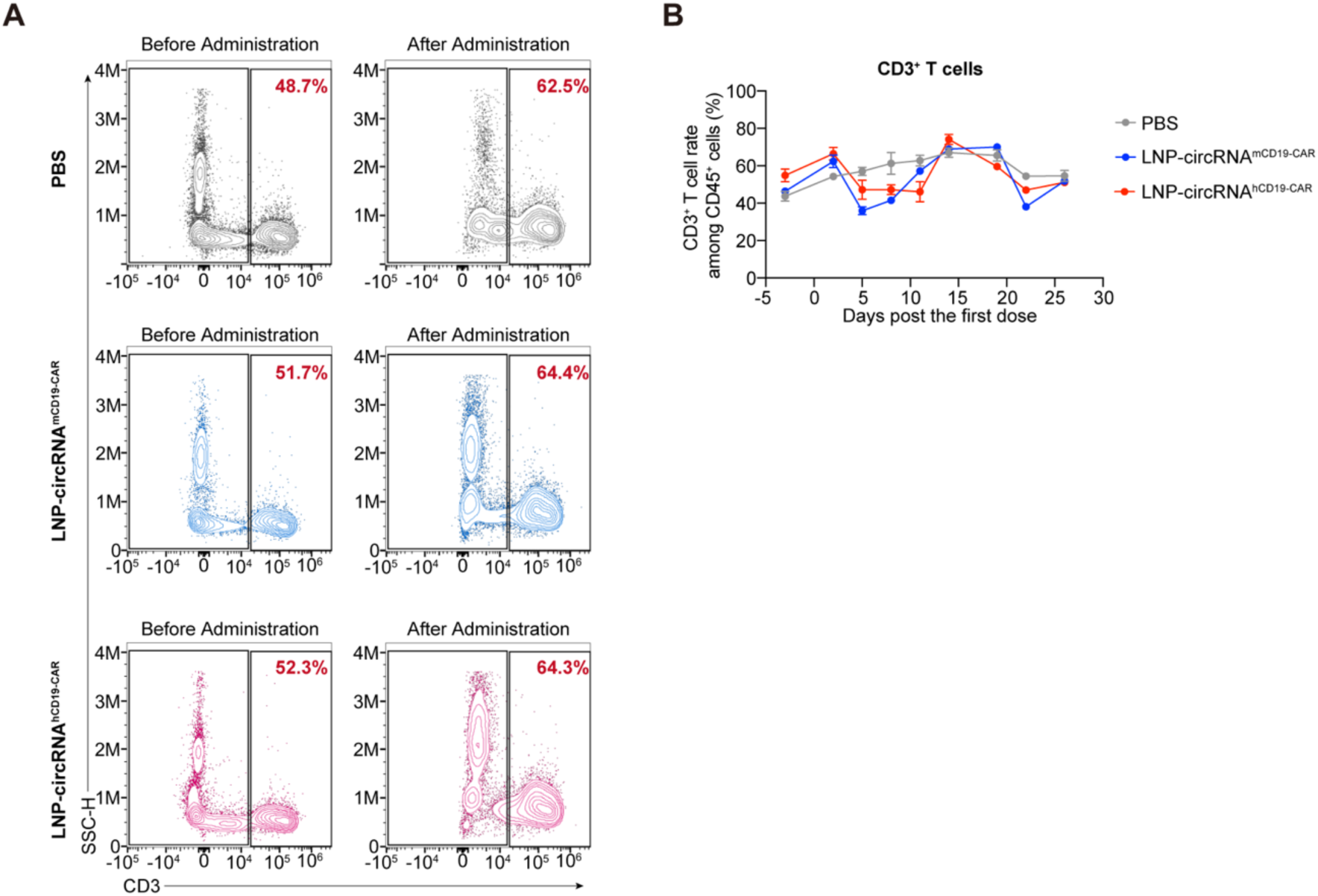
*In vivo* panCAR exhibited minimal effects on the CD3⁺ T cell populations of peripheral blood in MRL/MpJ-Fas^lpr^ mice, related to Figure 3. (A) Flow cytometry plots showing CD3⁺ T cells populations in peripheral blood of MRL/MpJ-Fas^lpr^ mice after PBS, circRNA^hCD19-CAR^ or circRNA^mCD19-CAR^ administration at 14 days post the first dose. (B) Effects of circRNA^hCD19-CAR^ or circRNA^mCD19-CAR^ on CD3⁺ T cells populations in peripheral blood of MRL/MpJ-Fas^lpr^ mouse models. In (B), data are presented as the mean ± SEM.

**Figure S5.**
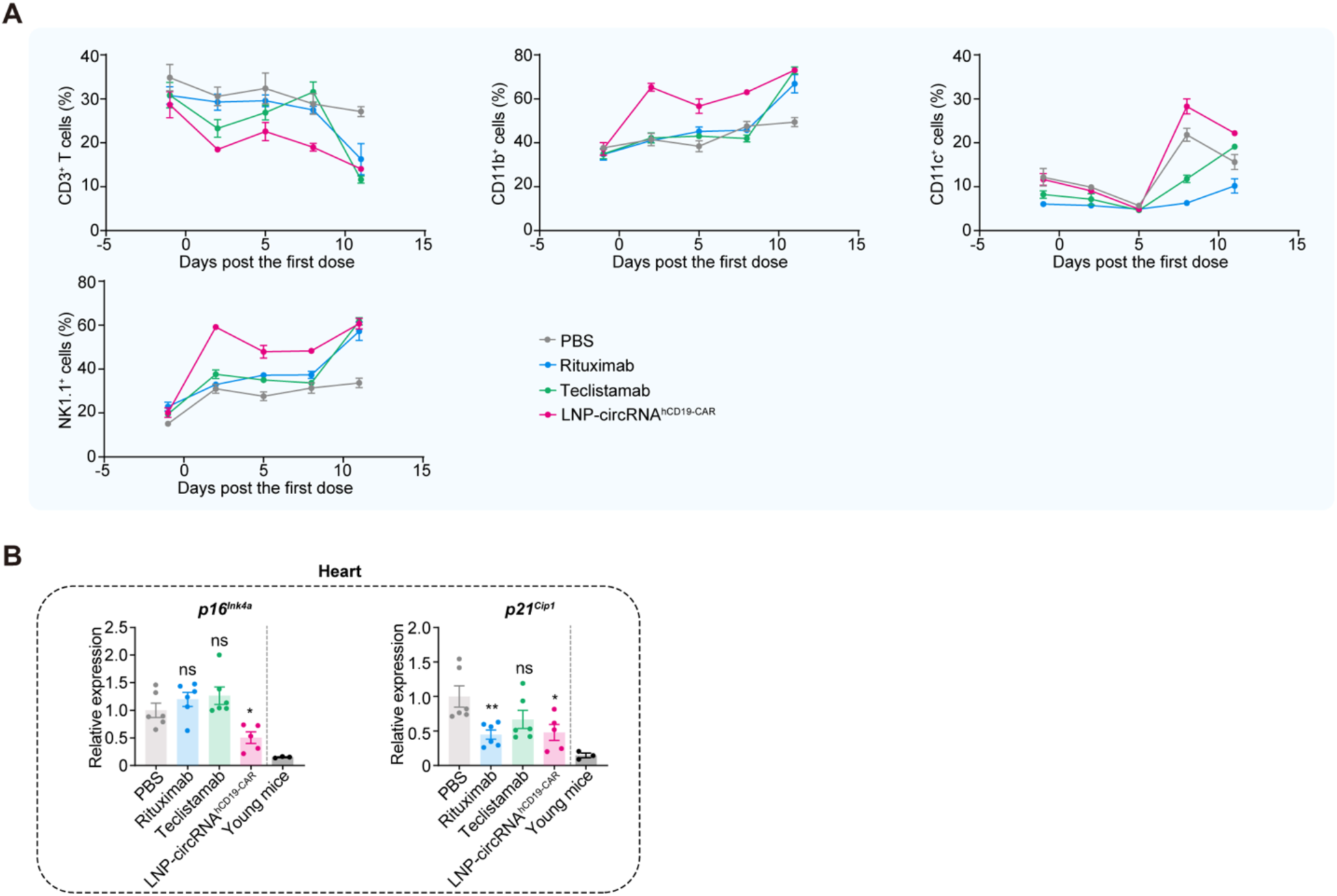
*In vivo* panCAR attenuated aging-associated phenotypes in 17-month-old aging mice, related to Figure 5. (A) Comparation of *in vivo* panCAR with antibody-based therapeutics on distinct immune cell populations in the peripheral blood of aging mice after treatments. (B) Comparative analysis of antibody or *in vivo* panCAR treatment on aging-associated gene expression in heart tissues using RT-qPCR. Each symbol represents an individual mouse. In (A) and (B), data are presented as the mean ± SEM. In (B), an unpaired two-sided Student’s *t* test was performed for comparison as indicated in the figures, \**p* < 0.05; \*\**p* < 0.01; \*\*\**p* < 0.001; \*\*\*\**p* < 0.0001; ns, not significant.

**Figure S6.**
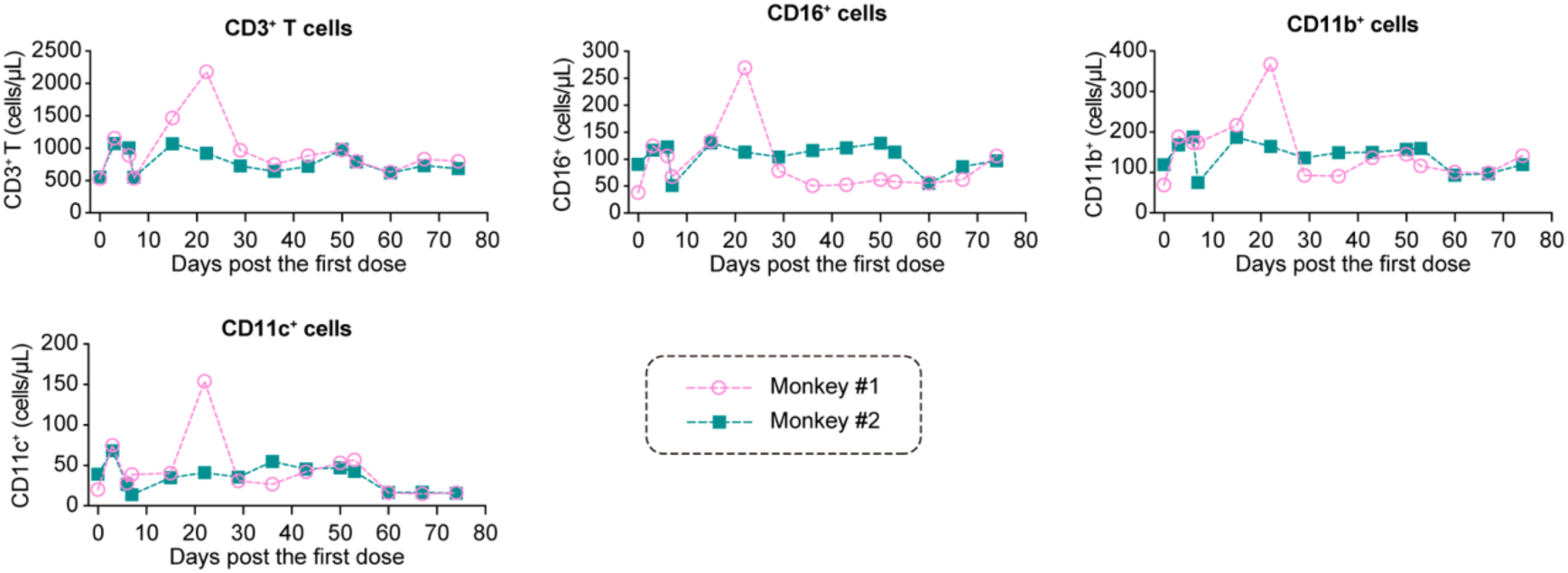
Effects of *in vivo* panCAR on T cells and other immune cell types in peripheral blood of cynomolgus monkey, related to Figure 6.

## RESOURCE AVAILABILITY

### Lead contact

Further information and requests for resources and reagents should be directed to and will be fulfilled by the lead contact, Liang Qu.

### Material availability

All unique reagents generated in this study are available from the lead contact (L.Q.) with a completed Material Transfer Agreement.

### Data and code availability

The data supporting the findings of this study are available from the lead contact (L.Q.) upon a reasonable request under a completed Material Transfer Agreement. This study did not generate any unique code. Any additional information required to re-analyze the data reported in this study is available from the lead contact upon request.

## ACKNOWLEDGMENTS

We acknowledge members of the Qu Laboratory for discussion throughout this study. We thank the Core Facility of Shanghai Medical College, Fudan University and the Public Technology Platform of School of Basic Medical Sciences, Fudan University for technical supports. This project was supported by funds from National Key R&D Program of China (2023YFC2604300 and 2024YFC3408200 to L.Q.); Shanghai Pilot Program for Basic Research - Fudan University 21TQ400100 (25TQ007 to L.Q.); Shanghai Rising-Star Program (24QA2701600 to L.Q.); Key Project in Synthetic Biology of Science and Technology Commission of Shanghai Municipality (23HC1400100 to L.Q.); Shanghai Municipal Health Commission Collaborative Innovation Cluster Project (2024CXJQ02 to L.Q.); Morning Glory Plan of Shanghai Municipal Education Commission (to L.Q.); Research Project Plan of Shanghai Municipal Health Commission (20234Y0235 to L.Q.).

## AUTHOR CONTRIBUTIONS

L.Q. conceived and supervised this project. L.Q., Y.W., X.W., Q.P., Y.Wu and T.Y. designed the experiments. Y.W., X.W., Q.P., H.K., C.Z., S.S., J.Y., X.L., W.W. and L.Z. performed the preparation of circRNAs and mRNAs, cell experiments, mouse experiments, detection experiments, and data collection with the help of L.Q. Y.W. and X.W. prepared the LNPs with the help of L.Q. Y.W., X.W., and Q.P. performed the monkey experiments and data collection with the help of T.Y., Y.Wu and L.Q. L.Q., Y.W., X.W. Q.P., Y.Wu, T.Y., H.X., H.K. and C.Z. wrote this manuscript with the help of other authors.

## METHODS

### Cell culture

The HEK293T cell line was sourced from our laboratory, while the THP-1, J774A.1 and Raji-Luc-EGFP were procured from Shanghai Zhong Qiao Xin Zhou Biotechnology Co., Ltd. The hCD19-overexpressing cancer cell line (A549-hCD19-EGFP) was procured from Vigen Biotechnology (Zhenjiang) Co., Ltd. THP-1 and Raji-Luc-EGFP were cultured in RPMI 1640 medium supplemented with 10% fetal bovine serum and 1% penicillin-streptomycin. HEK293T, J774A.1, and A549-EGFP-hCD19 cell lines were cultured in DMEM supplemented with 10% fetal bovine serum and 1% penicillin-streptomycin. All cell lines were maintained in a 5% CO2 incubator at 37°C and were confirmed to be mycoplasma-free. Human peripheral blood mononuclear cells (PBMCs) were used for *in vitro* killing experiments. Human PBMCs were isolated from a healthy male donor born in 1993. Subsequently, T cells were isolated using a T cell isolation kit and cultured in RPMI 1640 medium supplemented with human recombinant interleukin-2 (IL-2), human T-activator CD3/CD28 beads, 10% FBS, and 1% penicillin-streptomycin. The human primary T cells were randomly allocated into three groups: untreated, circRNA^Ctrl^, and circRNA^hCD19-CAR^, with each group containing 1 × 10^6^ cells for *in vitro* killing assays.

### CircRNA transfection *in vitro*

HEK293T cells were seeded in 24-well plates at a density of 1.5 × 10⁵ cells per well. 12 hours later, the cells were transfected with 2 μg circRNA or mRNA per well using Lipofectamine MessengerMax (Invitrogen, #LMRNA003) according to the manufacturer’s protocol. Cells were harvested 24 hours post-transfection for subsequent analysis. For THP-1 and J774A.1 cell lines, RNA transfection was performed via electroporation following the manufacturer’s instructions.

### Isolation and transfection of primary T Cells

Human Peripheral blood mononuclear cells were obtained from Shanghai Hycells Biotechnology Co., Ltd. T cells were subsequently isolated using a T cell isolation kit (STEMCELL, #17951). The isolated T cells were activated and expanded by adding recombinant human interleukin-2 (rhIL-2, MCE, #HY-P70758) and human T cell activator CD3/CD28 beads (Thermo, #11131D). After approximately one week of culture, circRNA^hCD19-CAR^ was transfected into the cells via electroporation. Cells were harvested 24 hours post-transfection, and anti-hCD19 CAR expression was assessed by flow cytometry.

### *In vitro* tumor cytotoxicity assay

A549-hCD19-EGFP cells were seeded in 96-well white plates at a density of 6 × 10^3^ cells per well and Raji-Luc-EGFP cells were seeded in 96-well white plates at a density of 2 × 10⁴ cells per well. Subsequently, circRNA^hCD19-CAR^ -transfected primary T cells or macrophages were added at various effector-to-target (E:T) ratios. For A549-hCD19-EGFP tumor cells, after a 48-hour incubation, rinse the effector cells with Hank’s buffer, followed by the addition of 100 μL fresh medium and 50 µL XTT reagent. The plate was then incubated at 37°C for 4 hours. Subsequently, absorbance was measured at 450 nm using an ELISA reader, with 650 nm as the reference wavelength. For Raji-Luc-EGFP tumor cells, following 48 h of co-culture at 37°C, luciferase activity was measured using BRITELITE PLUS kit (PerkinElmer).

### Animals and ethics statement

Female BALB/c mice (6- to 8-week old) were ordered from Beijing Vital River Laboratory Animal Technology Co., Ltd. Female aging BALB/c mice (17-month-old) were obtained from Biocytogen Pharmaceuticals (Beijing) Co., Ltd. The female MRL/MpJ-Fas^lpr^ were procured from Cyagen Biosciences Inc. Reporter BALB/c mice carrying the CAG-loxp-zsGreen-stop-loxp-tdTomato-PolyA construct were acquired from GemPharmatech Co., Ltd. All mice were maintained under specific pathogen-free (SPF) conditions in the Laboratory Animal Center of Fudan University. All animal experiments were approved by Fudan University Laboratory Animal Center (Shanghai) and conducted in accordance with the National Institute of Health Guide for Care and Use of Laboratory Animals.

### Expression of circRNA *in vivo*

To assess the efficiency of targeting different immune cells within secondary lymphoid organs, mice were first intravenously injected with 15 μg of circRNA^Cre^ encapsulated in two distinct LNPs. Forty-eight hours post-injection, spleens and lymph nodes were harvested from the reporter mice. The editing efficiency of circRNA^Cre^ in various immune cell populations was then determined via flow cytometry. Briefly, single-cell suspensions were prepared from the harvested spleens and lymph nodes. Cells were washed with staining buffer (PBS supplemented with 2% FBS), add the Live/Dead dye and incubate at room temperature (RT) for 20 min. After washing, cells were incubated with fluorochrome-conjugated antibodies targeting specific immune cell surface markers for 30 minutes at 4°C in the dark, washed three times with staining buffer, resuspended, and subsequently analyzed using flow cytometry.

To evaluate CAR expression levels in different immune cells *in vivo*, 6-8-week-old BALB/c mice were intravenously administered with circRNA^Ctrl^ or circRNA^hCD19-CAR^ encapsulated in LNP#2. Six hours post-injection, peripheral blood, spleen, and lymph nodes were collected from each treatment group. Single-cell suspensions were prepared and cells were washed once with staining buffer, add the Live/Dead dye and incubated at room temperature (RT) for 20 min. Subsequently, after washing once, surface-marker antibodies for distinct immune cell populations were added. Since the CAR molecule carries an N-terminal Flag tag, an anti-Flag flow cytometry antibody was simultaneously introduced to detect CAR expression. Cells were then incubated for 30 minutes at 4°C in the dark. After washed three times with staining buffer, cells were resuspended and analyzed via using Cytek Aurora full spectrum flow cytometry. Data analysis was conducted with FlowJo software.

### Extraction of tissue RNA and qPCR

Total RNA was extracted from tissues of mice in different treatment groups according to the manufacturer’s protocol (TIANGEN, #DP451). Briefly, tissues were ground in liquid nitrogen, and tissues (10 mg) were lysed with lysis buffer. Subsequently, 10 μL of Proteinase K was added, followed by incubation at room temperature for 5 minutes. Following centrifugation at 12,000 rpm for 5 minutes, an equal volume of 70% ethanol was added to the supernatant. The mixture was centrifuged, and the pellet treated with DNase I at room temperature for 15 minutes. Following two-time washes, RNA was eluted in solution. The isolated RNA was then used to prepare cDNA libraries. And Gene expression levels across different tissues and treatment groups were analyzed using the SYBR green fluorescent dye method, with *Gapdh* as the reference gene.

### Isolation and culture of bone marrow-derived macrophages (BMDMs)

The femurs and tibiae were dissected using sterile scissors. A 1 mL syringe was used to flush out the bone marrow from the femurs and tibiae with culture medium, repeating the flushing process 3–4 times until no visible red residue remained in the bones. The cell suspensions were then filtered through a 70-μm cell strainer into a 15 mL centrifuge tube and centrifuged at 1500 rpm for 5 minutes. The supernatant was discarded, and 1 mL of red blood cell lysis buffer was added. After incubation at room temperature for 5 minutes, PBS was added to a total volume of 10 mL to terminate lysis. The samples were centrifuged again at 1500 rpm for 5 minutes, followed by supernatant removal. Cells were resuspended in complete medium (DMEM supplemented with 10% FBS, 1% penicillin-streptomycin, and 20 ng/mL M-CSF), seeded in 6-well plate and cultured in a 37°C incubator. The medium was replaced every three days.

### Senescence-associated β-galactosidase (SA-β-Gal) staining

After one week of BMDMs culture, senescence-associated β-galactosidase staining was performed according to the manufacturer’s instructions of the senescence β-galactosidase staining kit. Briefly, the cell culture medium was discarded, and the cells were washed once with PBS. Then, 1 mL of β-galactosidase staining fixative solution was added per well, and the cells were fixed at room temperature for 15 minutes. After removing the fixative, the cells were washed three times with PBS. Subsequently, 1 mL of the staining working solution was added per well. The cells were incubated overnight at 37°C, protected from light and without CO₂. Finally, cells were observed and recorded under a standard optical microscope, and the number of positive cells was quantified using ImageJ.

### Flow cytometry analysis

For the cell lines, after collecting the transfected cells and washing once with staining buffer (PBS supplemented with 2% FBS), anti-Flag antibody was added at a 1:200 dilution. Following incubation at 4°C in the dark for thirty minutes, the cells were washed three times with staining buffer, resuspended, and circRNA expression was detected by flow cytometry.

For analysis of murine B cell depletion *in vivo*, following the acquisition of a single-cell suspension, the cells were washed once with staining buffer (PBS supplemented with 2% FBS). They were then stained with Live/Dead dye at room temperature for 20 minutes. After once wash, various cell surface antibodies were added, and the cells were incubated in the dark at 4°C for 30 minutes. Subsequently, the cells were washed three times with staining buffer and finally resuspended for analysis.

For the detection of B cell depletion in peripheral blood of cynomolgus monkeys, the following procedure was performed: Firstly, three volumes of red blood cell lysis buffer were added, and lysis proceeded for 10 minutes at room temperature. Then, eight volumes of PBS were added to stop the lysis. After centrifugation, cells were washed once with staining buffer (PBS supplemented with 2% FBS). Live/Dead dye was added, and cells were stained for 20 minutes at room temperature protected from light. Following another wash with staining buffer, the following surface antibodies were added: CD45 (BD, #558411), CD3 (BD, #552127), CD19(ABclonal, #A23008), CD20 (BioLegend, #302330), CD138 (BioLegend, #356520), CD11b (BioLegend, #101228), CD11C (BioLegend, #301638), CD16 (BioLegend, #302018), and Flag (BioLegend, #637330). Cells were incubated with the antibodies at 4°C for 30 minutes in the dark, followed by three washes. After resuspension, cells were filtered through a 70-μm cell strainer. For absolute quantification by flow cytometry, absolute counting beads were added according to the manufacturer’s instructions (Biolegend, #424902). Samples were then acquired on Cytek Aurora full spectrum flow cytometry, and data were analyzed using FlowJo software.

### Enzyme linked immunosorbent assay (ELISA) analysis

Serum samples were collected from cynomolgus monkeys at various time points. Serum biochemical parameters including IL-6 (JONLNBIO, #JL21801), IL-1β (Cloud-clone, #SEA563Si), TNF-α (Cloud-clone, #SEA133Si), MCP-1 (Cloud-clone, #SEA087Si), CRP (JONLNBIO, #JL21793), alanine aminotransferase (ALT) (COIBO, #CB10421-MK), aspartate aminotransferase (AST) (COIBO, #CB10220-MK), and alkaline phosphatase (ALP) (COIBO, #CB10488-MK) were analyzed according to the manufacturer’s instructions for the respective ELISA kits.

## QUANTIFICATION AND STATISTICAL ANALYSIS

The unpaired two-sided Student’s *t* test or paired Student’s *t* test was performed for comparison as indicated in the figure legends. Statistical analyses were performed with Prism 9.5 (GraphPad Software, Inc.).

